# Single cell analysis of gene expression in the substantia nigra pars compacta of a pesticide-induced mouse model of Parkinson’s disease

**DOI:** 10.1101/2022.02.18.481079

**Authors:** Arshad H. Khan, Lydia K. Lee, Desmond J. Smith

**Affiliations:** Department of Molecular and Medical Pharmacology, David Geffen School of Medicine, UCLA, Los Angeles, CA 90095-1735; Department of Obstetrics and Gynecology, David Geffen School of Medicine, UCLA, Los Angeles, CA 90095-6928

## Abstract

Exposure to pesticides in humans increases the risk of Parkinson’s disease (PD), but the mechanisms remain poorly understood. To elucidate these pathways, we dosed C57BL/6J mice with a combination of the pesticides maneb and paraquat (MNPQ). Behavioral analysis revealed motor deficits consistent with PD. Single cell RNA sequencing of substantia nigra pars compacta revealed both cell-type specific genes and genes expressed differentially between pesticide and control, including *Fam241b, Emx2os, Bivm, Gm1439, Prdm15* and *Rai2*. Neurons had the largest number of significant differentially expressed genes, but comparable numbers were found in astrocytes and less so in oligodendrocytes. In addition, network analysis revealed enrichment in functions related to the extracellular matrix. These findings emphasize the importance of support cells in pesticide-induced PD and refocus our attention away from neurons as the sole agent of this disorder.

## Introduction

Parkinson’s disease (PD) ranks as the second most common neurodegenerative disorder (Kalia and Lang, 2015). The disease affects approximately 2 to 3% of the world population over 65 years of age and the risk of PD increases with age. In the US alone, more than one million people have PD and about 50,000 new cases are diagnosed yearly. The cardinal signs of the disease are tremor, rigidity, bradykinesia and postural instability. There are, in addition, a wide variety of other features including cognitive dysfunction, dementia, mood disorders, autonomic dysfunction and olfactory disturbances. Pathologically, PD is characterized by loss of dopaminergic neurons in the substantia nigra pars compacta (SNpc) that project to medium spiny neurons in the dorsal striatum (Fahn and Sulzer, 2004; Zhong et al., 2021). Surviving neurons in the SNpc exhibit characteristic inclusions called Lewy bodies, which are comprised principally of α-synuclein protein (Fernagut et al., 2007).

Genetics makes a substantial contribution to PD. The heritability of PD due to common variants is ∼22%, and 90 such variants have been identified (Nalls et al., 2019). In addition, more than 20 monogenic loci have been uncovered (Blauwendraat et al., 2020). However, environment clearly also makes an important contribution to PD. Epidemiological studies have shown that exposure to pesticides increases the risk of PD (Chambers-Richards et al., 2021; Fernagut et al., 2007; Hatcher et al., 2008). In particular, both the fungicide maneb (MN, manganese ethylene-bis-dithiocarbamate) and the herbicide paraquat (PQ) have been associated with PD (Ascherio et al., 2006; Wang et al., 2011).

In mice, either MN or PQ results in neurodegeneration of dopaminergic neurons by inhibiting mitochondrial function and elevating oxidative stress, a common pathway for PD (Chin et al., 2008; Helley et al., 2017; Smith, 2009; Wang et al., 2011). A well-established mouse model of pesticide-induced PD employs combined dosing of both agents (MNPQ). The result is loss of tyrosine hydroxylase neurons in the substantia nigra pars compacta (SNpc), increased α-synuclein aggregates and abnormalities in motor behavior reminiscent of PD in humans (Fernagut et al., 2007; Gollamudi et al., 2012; Richter et al., 2017; Wang et al., 2011).

Transcript profiling of human PD brain samples and animal models have revealed molecular pathways underpinning the disorder. As well as disruption of mitochondrial function and oxidative stress, these pathways include dopamine metabolism, protein degradation, inflammation, vesicular transport and synaptic transmission (Borrageiro et al., 2018; Brown et al., 2002; Chin et al., 2008; Greene, 2012). More recently, massively parallel single cell RNA sequencing (scRNA-seq) has deciphered pathways of PD at finer cellular resolution (Ma and Lim, 2021). For example, studies using mouse and human samples have identified specific gene expression changes not only in dopaminergic neurons but also in oligodendrocytes, though not microglia (Agarwal et al., 2020; Bryois et al., 2020; Errea and Rodriguez-Oroz, 2021; Skene and Grant, 2016). One investigation found early downregulation of HDAC4-controlled genes in an induced pluripotent stem cell (iPSC) model of PD (Lang et al., 2019).

Despite the increasing use of scRNA-seq to understand PD, this technology has been little employed to decipher the cellular heterogeneity of pesticide-induced PD. In this report, we evaluate the cellular and gene expression changes occurring in the SNpc in a mouse model of PD induced using MNPQ.

## Materials and Methods

### Pesticide treatment

A total of thirty-six C57BL/6J mouse (8 weeks old) were obtained from Jackson Laboratory, Bar Harbor, Maine. Each mouse was housed for two weeks (3 per each cage) to allow acclimatization to the new environment. Mice were treated either with saline (vehicle) or MNPQ (9 males, 9 females in each group). Animals were weighed at 10 weeks of age and intraperitoneal (i.p.) injections of 10 mg kg^-1^ PQ and 30 mg kg^-1^ MN given twice per week (Monday and Friday) for three weeks. PQ was administered first, followed an hour later by MN. Control mice received saline under the same regimen. Two females in the MNPQ group died before data could be collected.

### Pole and adhesive (dot) removal test

One week after administration of MNPQ or saline, motor effects were evaluated in treated and control mice using the pole test and the adhesive (dot) removal test (Fernagut et al., 2007; Taylor et al., 2009). For the pole test, each mouse was placed head-up on top of a vertical wooden pole with a rough surface, 50 cm in height and 1 cm in diameter. The animals were allowed to orient themselves downward and to descend along the pole back into their home cage. Each mouse was exposed to three trials, and the time spent to orient downward (t-turn) and the time to descend (t-descend) were recorded. If the mouse was unable to turn downward, the default value of 120 s was recorded as the maximal severity of impairment. For the adhesive (dot) removal test, each mouse was removed from their home cage and placed them in a testing cage for 60 seconds. After acclimatization, a 1.3 cm diameter adhesive paper was placed on top of their forehead. Each mouse was given three trials, and the times to touch the dot and to remove the dot were recorded.

Linear mixed models were used to analyze the pole and adhesive (dot) removal test, with fixed effects of treatment and sex, and a random intercept of individual mouse. Significance testing of the fixed effects used a t-statistic with Kenward-Roger degrees of freedom. Measurements are quoted as means ± standard error of the mean (s.e.m.)

### Single cell isolation from SNpc

One week after completion of behavioral testing, three males from the control (vehicle) and three males from the treated (MNPQ) mice were used for single cell isolation from SNpc. Animals were euthanized using isoflurane followed by cervical dislocation. Immediately after euthanasia, the brains were removed and placed on an ice cold mouse brain matrix (1 mm slices) and the region of 1.28 mm bregma to 2.28 mm bregma containing the SNpc sliced out. Punch dissection was used to dissect out SNpc and single cells dissociated by digestion using 2 mg ml^-1^ papain for 30 min at 34βC followed by trituration for 35 minutes. Debris were removed by filtration using a 40 mm filter. Cells were pelleted and supernatant removed. Cells were resuspended in 1 × PBS with 0.04 % FBS and quantified and quality checked using a Countess II Automated Cell Counter (Thermo Fisher). A total of 1 ml of cells at ∼1200 cells μl^-1^ were submitted for sequencing.

### Single cell RNA sequencing

Sequencing libraries were constructed from isolated single cells of vehicle and MNPQ samples using 10X Genomics Chromium technology with 3’ end gene expression library preparation. An Illumina NextSeq 500 SBS sequencing machine was used with 1 × 75 cycles and paired end sequences of 26 bp and 57 bp. Sequence data were demultiplexed and mapped against the indexed mouse reference genome (refdata-cellranger-mm10-3.0.0.tar.gz; GRCm38/mm10) using the Cell Ranger count pipeline software package (10X Genomics). Data from each sample were initially filtered to exclude genes expressed in fewer than five cells, and to exclude cells that contained fewer than 100 expressed genes.

To improve quality, strict filtering was further done on the raw data using the Seurat R package with the number of unique molecular identifiers (nUMI) > 500, nGene > 500, log_10_GenesPerUMI > 0.80 and mitoRatio < 0.20 criteria (Hao et al., 2021). To account for differences in sequencing depth per cell for each sample, normalization and variance stabilization of the scRNA-seq data was done using the SC transform method in Seurat.

### Cell clustering and identification of marker genes

Cell types in each sample were clustered using dimensionality reduction procedures, such as principal component analysis (PCA) and uniform manifold approximation and projection (uMAP). Graph-based clustering was performed using the Seurat function FindNeighbors and FindClusters. Each cluster was separated using the Leuvain algorithm with a resolution parameter of 0.5. To visualize the clusters, non-linear dimensional reduction was performed using uMAP with the same principal components employed for the graph-based clustering. The Seurat function FindAllMarkers was used to identify marker genes for the clusters.

### Average expression and differential expression of each marker

The average expression of cell type specific markers was calculated using the Seurat package. Markers for each cell types were subset and natural logarithms were taken of average expression of RNA counts plus one. Differential expression of genes and corresponding significance values between control and experimental samples were calculated separately for each cluster using the FindMarkers functions of Seurat package.

### Data availability

The sequencing data generated in this study can be downloaded from the NCBI BioProject database (https://www.ncbi.nlm.nih.gov/bioproject/) under accession number PRJNA756542. Data also available from figshare (https://figshare.com/; doi: 10.6084/m9.figshare.19197056)

## Results

### Pole and adhesive removal test

On the pole test, the MNPQ treated mice showed significantly greater time to turn around (control mice = 5.0 ± 0.37 s, MNPQ mice = 6.5 ± 0.40 s, t[1,31] = 2.7, *P* = 0.012) and also to climb down the pole (control mice = 11.4 ± 0.58 s, MNPQ mice = 13.7 ± 0.62 s, t[1,31] = 2.8, *P* = 9.5 × 10^−3^) (Figure 1A,B). There was no significant effect of sex for either the turn around or climb down time.

**Figure 1.**
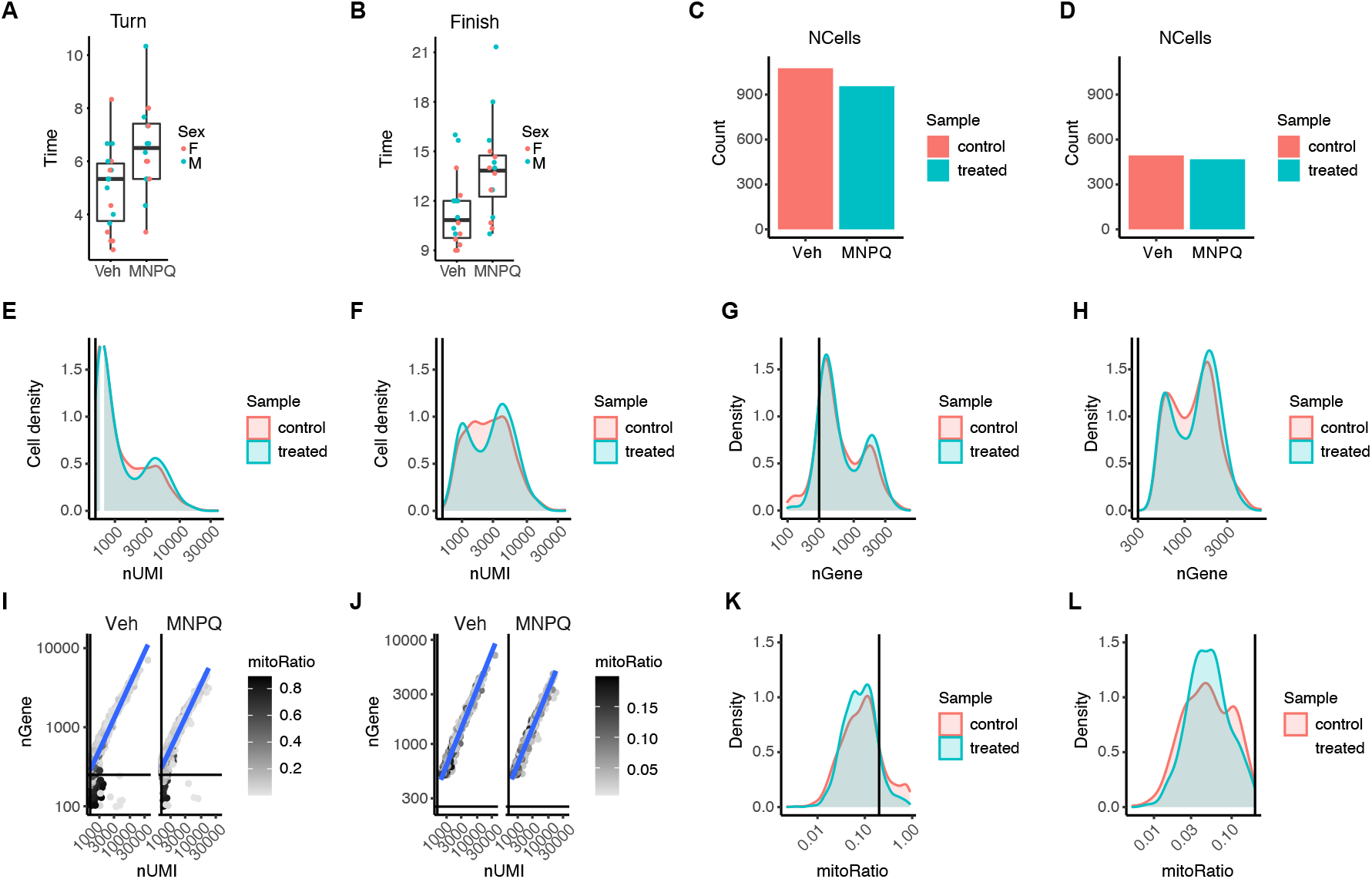
Behavioral testing and data filtering methods. (**A**) Pole test, time to turn. (**B**) Pole test, time to finish climbing down pole. Veh, vehicle; MNPQ, maneb and paraquat. Differences between groups for both endpoints, P < 0.05. (**C**) Cell counts using Cell Ranger pipeline. (**D**) Cell counts using strict filtering removing unwanted cells that are either dead or tagged with ambient RNA. (**E**) Number of transcripts from Cell Ranger. (**F**) Number of transcripts from strict filtering. (**G**) Number of genes per cell obtained by Cell Ranger. (**H**) Number of genes per cell obtained by strict filtering criteria. (**I**) High mitochondrial read fractions (dark color) indicate damaged/dead cells due to leaked cytoplasmic mRNA. (**J**) Mostly live cells after filtering out dead cells. (**K**) Appreciable numbers of dead cells before strict filtering using mitochondrial read fractions. (**L**) Dead cells removed after strict filtering.

For the adhesive removal test, there was no significant difference between vehicle and MNPQ treated mice for time to either touch the dot (control mice = 11.7 ± 1.91 s, MNPQ mice = 15.9 ± 2.03 s, t[1,31] = 1.52, *P* = 0.14) or for time to remove the dot (control mice = 13.3 ± 1.99 s, MNPQ mice = 17.9 ± 2.11 s, t[1, 31] = 1.58, *P* =0.12). There was no significant effect of sex for either the time to touch or remove the adhesive dot. Nevertheless, the pole test was consistent with the notion that pesticide exposure in mice results in motor deficits reminiscent of PD.

### Single cell RNA sequencing and mapping

Sequencing results are summarized in Supplementary Table S1. More than 180 and 175 million sequence reads, 1105 and 965 cells, and 17480 and 16908 transcripts were obtained from SNpc of control and MNPQ animals, respectively. The sequencing data showed acceptable quality based on the percentage of bases with Q30 or better in the RNA reads, with scores of 64.5% and 63.8% in control and MNPQ samples (ideal threshold is > 65%). However, the fraction of reads in cells (the fraction of confidently mapped reads with cell-associated barcodes) was 24.7% and 23.9% for the control and experimental samples (ideal score of > 70%).

The poor-quality score for the fraction of reads in cells suggested that many of the reads were not assigned to cell-associated barcodes. Possible causes are high levels of ambient RNA or increased number of cells with low RNA content, preventing the algorithm from calling cells. Ambient RNAs are usually pooled mRNA molecules that have been released in the cell suspension from stressed cells or cells that have undergone apoptosis. Isolation of single cells from mouse SNpc is a tedious process, resulting in many dead cells. Despite the marginal quality of the data, we decided to pursue further analysis in the spirit of an exploratory study.

### Data filtering

We initially filtered the sequence data from each sample using the Cell Ranger pipeline to exclude genes expressed in fewer than five cells and cells containing fewer than 100 expressed genes. However, because of the poor quality of the scRNA-seq data, we decided to impose stricter thresholding than is standard by using the Seurat pipeline. The mitochondrial transcript ratio of each cell was incorporated into the metadata, thus avoiding over representation of mitochondrial genes that possibly represent dead cells (Achim et al., 2015; Macosko et al., 2015). We used the following criteria: number of unique molecular identifiers (nUMI) > 500, nGene > 500, log10GenesPerUMI > 0.80 and mitoRatio < 0.20.

We compared the cell counts per sample, UMI counts (transcripts) per cell and genes detected per sample after filtering from the Cell Ranger pipeline or using the strict filtering of the raw data from Seurat (Figures 1C-1H). We lost more than half of the cell number as a result of the strict filtering. Using the Cell Ranger filtering criteria, many cells had a UMI count < 500, while the strict filtering criteria selected cells with a UMI count > 500 (Figures 1E,F). A similar pattern was obtained from the number of genes detected per cell (Figures 1G,H). The distributions of UMI and genes per cell were bimodal, whereas a single peak is expected (Figures 1E-1H). The bimodal distributions may indicate the presence of biologically different cell types in the data (simpler vs more complex expression profiles) or cells that are larger in size (McGinnis et al., 2019; Shalek et al., 2013).

To determine whether the strong presence of cells with low numbers of genes/UMIs were due to mitochondria, we plotted the number of UMIs and number of genes detected per cell using the Cell Ranger pipeline (Figures 1I,K). The bottom left quadrants of the plots in Figure 1I represent poor quality cells with a low number of genes and UMIs. High mitochondrial read fractions (dark color) were found in these cells and are probably indicative of damaged/dying cells which only retain mitochondria mRNA and have leaked cytoplasmic mRNA. The strict filtering (< 0.2 mitochondrial ratio) permitted selection of data with very low numbers of dying cells. (Figures 1J,L).

The stringent filtering allowed us to remove most of the noise from the data, although almost half of the filtered data (specifically cell numbers) generated by Cell Ranger were lost. The total number of cells dropped to 494 and 468 for control and experimental samples, respectively, and the number of genes dropped to 13,038. However, the strict quality control provided us with more confidence in the downstream analysis.

### Cell clustering

Initial clustering of data from control and treated samples resulted in thirteen distinct cell clusters (Supplementary Figure S1). To identify cell types in each cluster, the top 25 marker genes in each cluster were assessed for cell specificity using a database of cell markers in human and mouse (Zhang et al., 2019). If 50% or more of the top 25 marker genes belonged to a specific cell type, then the cluster was assigned that cell type. This process reduced the number of clusters to seven (astrocytes, endothelial cells, neuron, microglia, oligodendrocytes, mural cells and ependymal cells) (Figure 2).

**Figure 2.**
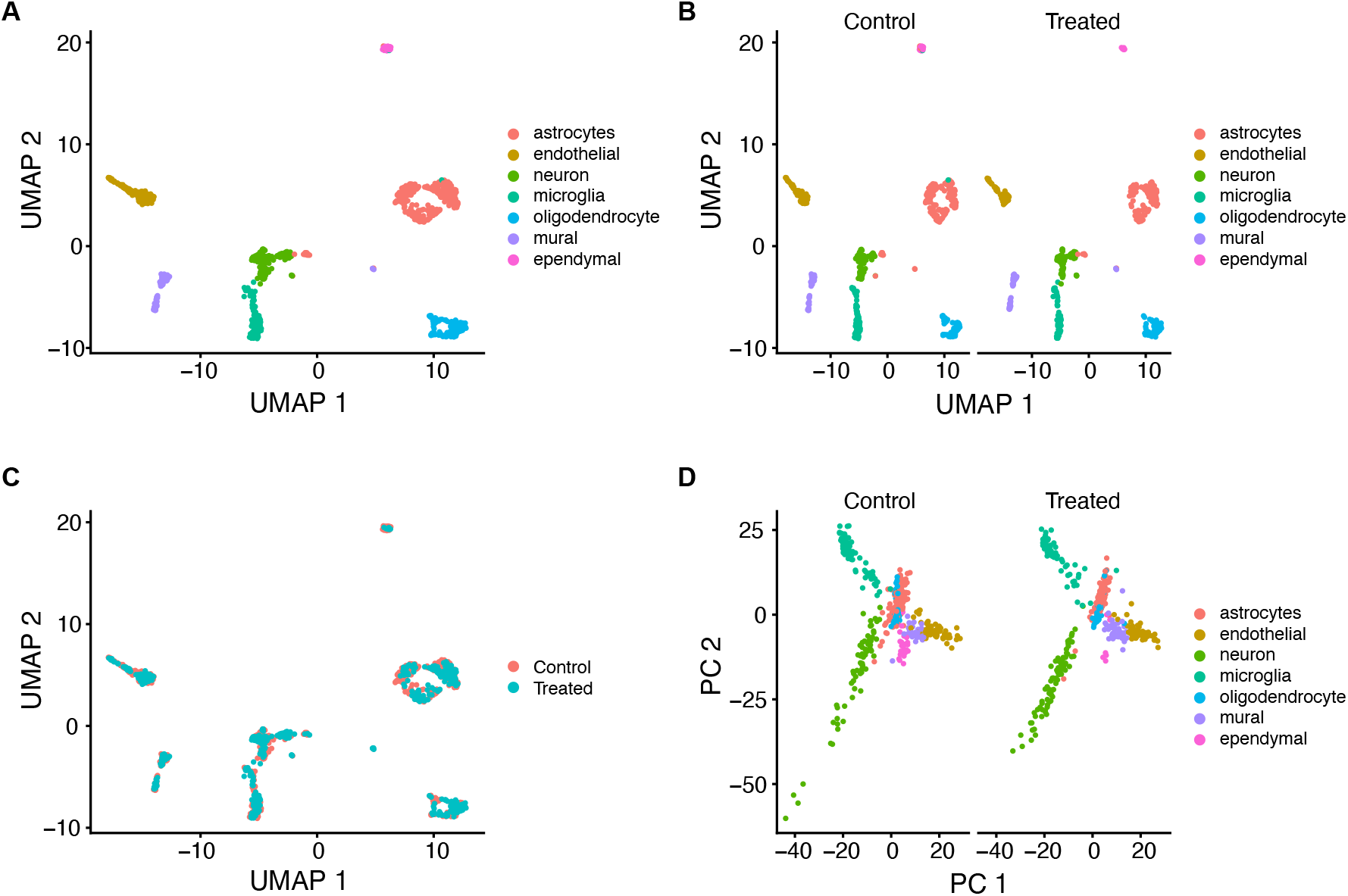
Cell clusters in SNpc from control and MNPQ mice. (**A**) Seven clusters classified by cell type using uniform manifold approximation and projection (UMAP). (**B**) Clusters separated by sample condition. (**C**) Overlap of cell clusters between samples. (**D**) Clusters separated by sample condition using principal components (PC).

Significant genes in each cell cluster based on expression differences with all other clusters are shown in Supplementary Table S2. Most of the marker genes were specific for their own cluster except for a few that overlapped. Examples of marker genes for each cell type in the combined samples and the separate MNPQ and control samples are shown using dot plots in Figure 3A,B. Heatmaps of mean expression for all markers in control and treated samples also showed cell type specific gene expression (Figure 3C,D).

**Figure 3.**
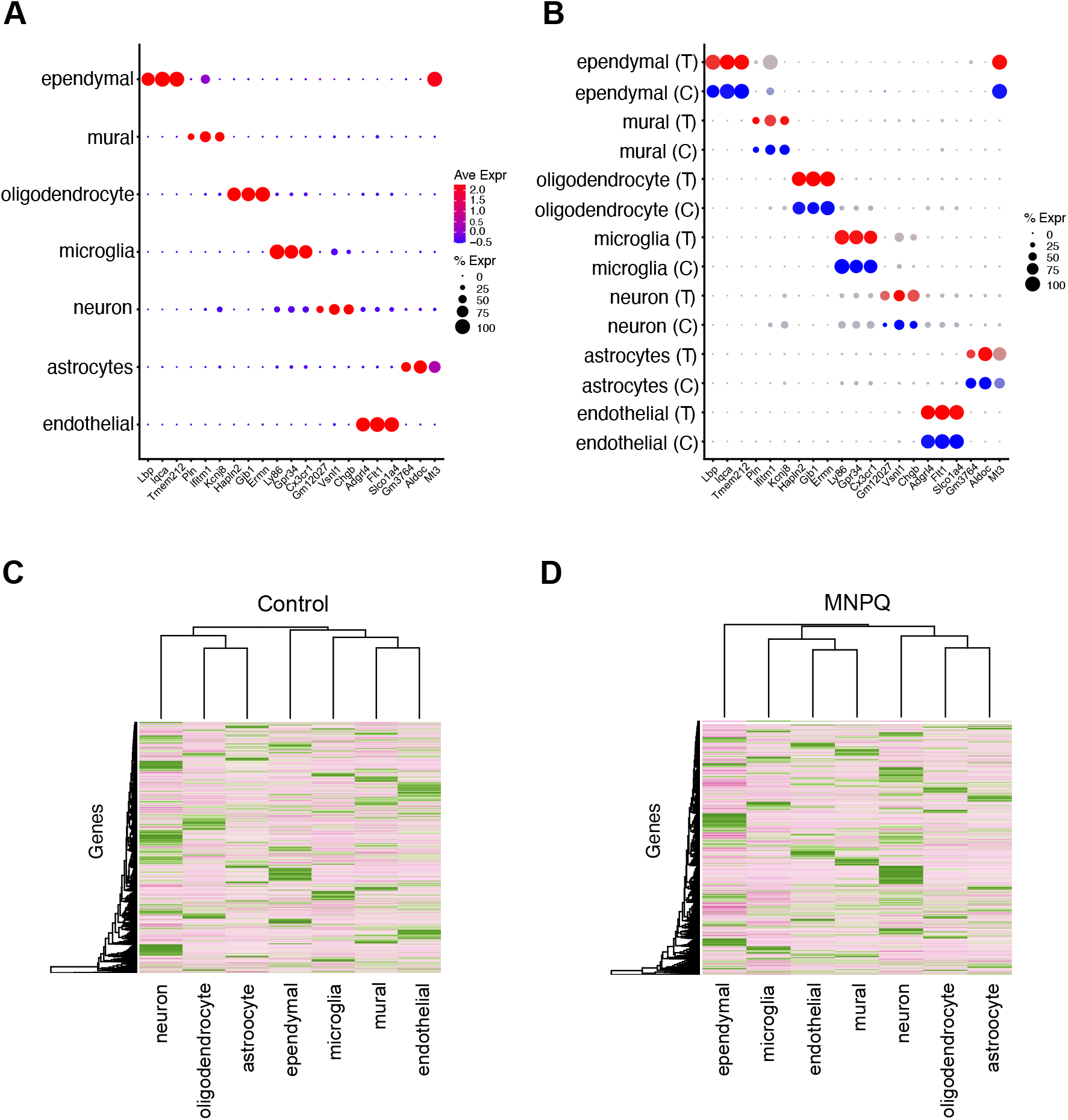
Marker genes for each cell cluster. (**A**) Marker genes for combined MNPQ and control samples. Average expression (Ave Expr) denoted by red-blue color scale. Percent of cells expressing gene (% Expr) denoted by dot size. (**B**) Same marker genes separated by sample type; MNPQ treated (T, red-grey expression scale) and control (C, blue-grey expression scale). (**C**) Heatmap for average expression of all control markers. (**D**) MNPQ samples. Cluster specific marker expression apparent.

### Gene markers in clusters

The top three most significant neuronal markers were *Syt1* (*P* = 1.91 × 10^−56^, 4.86 × 10^−58^), *Snap25* (*P* = 1.50 × 10^−49^, 2.72 × 10^−47^) and *Rtn1* (*P* = 2.90 × 10^−47^ and 9.13 × 10^−42^) (adjusted *P* values in MNPQ and control samples, respectively). The *Syt1* protein is thought to interact with α-synuclein and the *Snap25* protein forms part of the SNARE complex(Noor and Zahid, 2017). Changes in the abundance and distribution of the SNARE complex impairs dopamine-mediated modulation of synaptic function and is involved in PD initiation. *Rtn1* plays a role in neuronal injury in an *in vitro* model of PD using the neurotoxin MPP^+^ (1-methyl-4-phenylpyridinium) (Chang et al., 2019). Synaptophysin (*Syp*) was another strongly expressed neuronal gene in our study (adjusted *P* = 2.19 × 10^−46^ and 1.32 × 10^−30^ in MNPQ and control samples, respectively) and is a marker of synaptic terminals that shows loss in neurodegenerative disorders such as PD (Bai and Strong, 2014).

Astrocyte specific genes included *Gpr37L1* (*P* = 1.68 × 10^−70^ and 1.35 × 10^−59^), *Pla2g7* (*P* = 8.94 × 10^−63^ and 2.63 × 10^−52^) and *Prdx6* (*P* = 1.01 × 10^−48^ and 4.01 × 10^−32^) (adjusted *P* values in MNPQ and control samples, respectively). The protein encoded by *Gpr37L1*, the top astrocyte specific marker, is a G protein coupled receptor. Both the *Gpr37L1* protein and its homolog, the PD associated orphan receptor *Gpr37*, physically interact with the dopamine 2 receptor (*Drd2*), which is expressed in astrocytes as well as neurons (Hertz et al., 2019; Morató et al., 2021; Zhu et al., 2020). The *Gpr37* protein is also a key substrate for *Parkin*, mutations of which cause autosomal recessive juvenile PD (Imai et al., 2001).

Mutations in the astrocyte specific gene, *Pla2g7*, cause early onset PD (Magrinelli et al., 2021). In addition, transgenic mice expressing the astrocyte specific gene, *Prdx6*, show increased loss of dopaminergic neurons and more severe behavioral deficits in the MPTP (1-methyl-4-phenyl-1,2,3,6-tetrahydropyridine) mouse model of PD compared to non-transgenic controls (Yun et al., 2015).

The top oligodendrocyte specific markers were *Ermn* (*P* = 1.25 × 10^−88^ and 8.87 × 10^−87^), *Cldn11* (*P* = 4.70 × 10^−86^ and 3.50 × 10^−88^) and *Ugt8a* (*P* = 1.08 × 10^−73^ and 4.33 × 10^−88^) (adjusted *P* values in MNPQ and control samples, respectively). The oligodendrocyte marker *Hapln2* (adjusted *P* = 2.96 × 10^−66^ and 2.18 × 10^−79^ in MNPQ and control samples, respectively) promotes α-synuclein aggregation and may contribute to neurodegeneration in PD (Wang et al., 2016, 2019).

*C1qc*, which encodes complement subcomponent C1q, was specifically expressed in microglia (adjusted *P* = 3.09 × 10^−75^ and 4.82 × 10^−87^ in MNPQ and control samples, respectively). A meta-analysis of transcriptome data showed that this gene was more strongly expressed in the substantia nigra of PD patients compared to controls (Mariani et al., 2016). We also found high expression of *Meig1* in ependymal cells (adjusted *P* = 1.08 × 10^−42^ and 5.93 × 10^−86^ in MNPQ and control samples, respectively). The *Meig1* protein binds to the protein encoded by *Pacrg*, a gene co-regulated with *Parkin* (Khan et al., 2021).

*Cldn5* is essential for blood brain barrier (BBB) integrity (Ahn et al., 2021) and we found that this gene was differentially expressed in endothelial cells (adjusted *P* = 2.69 × 10^−76^ and 3.38 × 10^−88^ in MNPQ and control samples, respectively). Previous work has shown breakdown of the BBB in various neurological disorders, including PD (Obermeier et al., 2013).

### Functional enrichment of clusters

The top 10 marker genes showing the highest specificity in each cluster based on minimum *P* values were used to perform gene set enrichment analysis (GSEA) for biological process using g:Profiler (all adjusted *P* < 0.05; Supplementary Table S3) (Raudvere et al., 2019; Subramanian et al., 2005). Neurons revealed significant enrichments in calcium ion-regulated exocytosis of neurotransmitter, synaptic vesicle fusion to presynaptic active zone membrane, synaptic vesicle exocytosis, signal release from synapse and neurotransmitter secretion.

GSEA of biological process in astrocytes showed significant enrichment in positive regulation of neurofibrillary tangle assembly, inclusion body assembly and low-density lipoprotein particle remodeling, together with negative regulation of amyloid fibril formation. These biological processes are involved in Alzheimer’s disease, which has mechanistic overlaps with PD (Wakasugi and Hanakawa, 2021). In oligodendrocytes, enrichment was observed in ensheathment of neurons and axon, central nervous system myelination, oligodendrocyte differentiation and glial cell development.

GSEA of molecular function in neurons showed enrichment in syntaxin binding, SNARE binding and metal ion transmembrane transporter activity (Supplementary Table S3). Astrocytes showed enrichment in low-density lipoprotein particle receptor binding and tau protein binding.

### Expression differences between MNPQ and control mice

Genes with strong cluster specific expression and large expression differences between MNPQ and vehicle mice were identified (Supplementary Figures S2, S3 and S4, Supplementary Table S4). Relatively cluster enriched genes that showed large differences between MNPQ and control included *Dusp3, Pianp* (neurons), *Ppp1r3g, Tagln3* (astrocytes) and *Glul, Bsg* (oligodendrocytes). Some had known links to PD, including *Acp2, Rac1, Sgta* (neurons) and *Cntfr, Fabp5* (astrocytes) (Ashtari et al., 2016; Guiler et al., 2021; Kubota et al., 2021; Pöyhönen et al., 2019; Wang et al., 2021).

In addition to cluster specific differential expression of genes, there were also suggestions of cluster agnostic differential expression. For example, *Gm42418* was strongly downregulated in MNPQ compared to vehicle in all seven cell types. *Gm42418* is a lncRNA and the role of these enigmatic transcripts in brain function, and in neurological disorders such as PD, is becoming more widely appreciated (Andersen and Lim, 2018; Taghizadeh et al., 2021). Other differentially expressed genes shared between at least two cell types included *Actb, Acls3, Agt, Atp1b1, Btbd17, Camk2n1, Cmbl, Dbp, Fabp5, Fam23b, Id4, Igfbp2, Malat1, Mt2, mt-Nd4l, Nnat, Ndufb5, Nap1l3, Prpf4b, Rgs22, Rps26, Rpl28, Slc6a11* and *Tagln. Malat1* was downregulated by MNPQ in astrocytes and endothelial cells and is a lncRNA that appears to play a role in PD and other neurodegenerative disorders (Lu et al., 2020).

### Significant differential expression between MNPQ and control samples

*P* values for differential expression between MNPQ and controls in each cluster were evaluated. All cell clusters, except ependymal cells, had genes whose expression was significantly different between control and sample, giving a total of 1750 significant genes (Bonferroni adjusted *P* < 0.05) (Supplementary Table S5). Most genes were expressed at higher levels in the controls; less than one fifth (304) of the 1750 significant markers were upregulated in the MNPQ samples. The top three most significant genes in each cell type were *Fam241b, Nlk, Ss18l1* (astrocyte), *Emx2os, Tro, Ppip5k1* (endothelial cells), *Bivm, Cntrob, Nkrf* (microglia), *Gm1439, Samd15, Pdia5* (mural cells), *Prdm15, Sema3e, Capn15* (neuron), *Rai2, Dcbld1, Entpd7* (oligodendrocyte) (adjusted *P* < 1.81 × 10^−9^).

Among the statistically significant genes, 17 overlapped with genes for PD identified from genome-wide association studies (*P* < 5 × 10^−8^) (Buniello et al., 2019). These genes included *Fyn, Gak, Rit2*, and *Dgkq*. One gene, *Lrrk2*, is also implicated in monogenic PD. Neurons had the largest number of significant genes (472) followed by astrocytes (365), oligodendrocytes (266), endothelial cells (224), microglia (218) and mural cells (211). Our finding of abundant differentially expressed genes in astrocytes is consistent with the finding that this cell type plays an active role in dopaminergic signaling (Corkrum et al., 2020). The significant genes in oligodendrocytes echoes a recent scRNA-seq study of normal human substantia nigra suggesting a link between PD risk and these cells (Agarwal et al., 2020; Errea and Rodriguez-Oroz, 2021). Almost one third of significant differentially expressed genes were shared between two or more clusters. A total of 14 genes, including *Epm2a, Sgtb, 9330182L06Rik* and *Fam160b2*, overlapped in 4 or more clusters.

Genes with statistically significant differences in expression between MNPQ and vehicle in each cell type were illustrated in feature plots (Supplementary Figure S5). Selected genes in each cell type are shown in Figure 4A. Although MNPQ causes decreased average expression of most genes compared to vehicle, the percentage of cells with detectable expression was increased. Figure 4B,C shows heatmaps of the percent of cells expressing genes for all genes and for statistically significant differentially expressed genes. The increased numbers of cells expressing statistically significant but down-regulated genes in the MNPQ samples is apparent.

**Figure 4.**
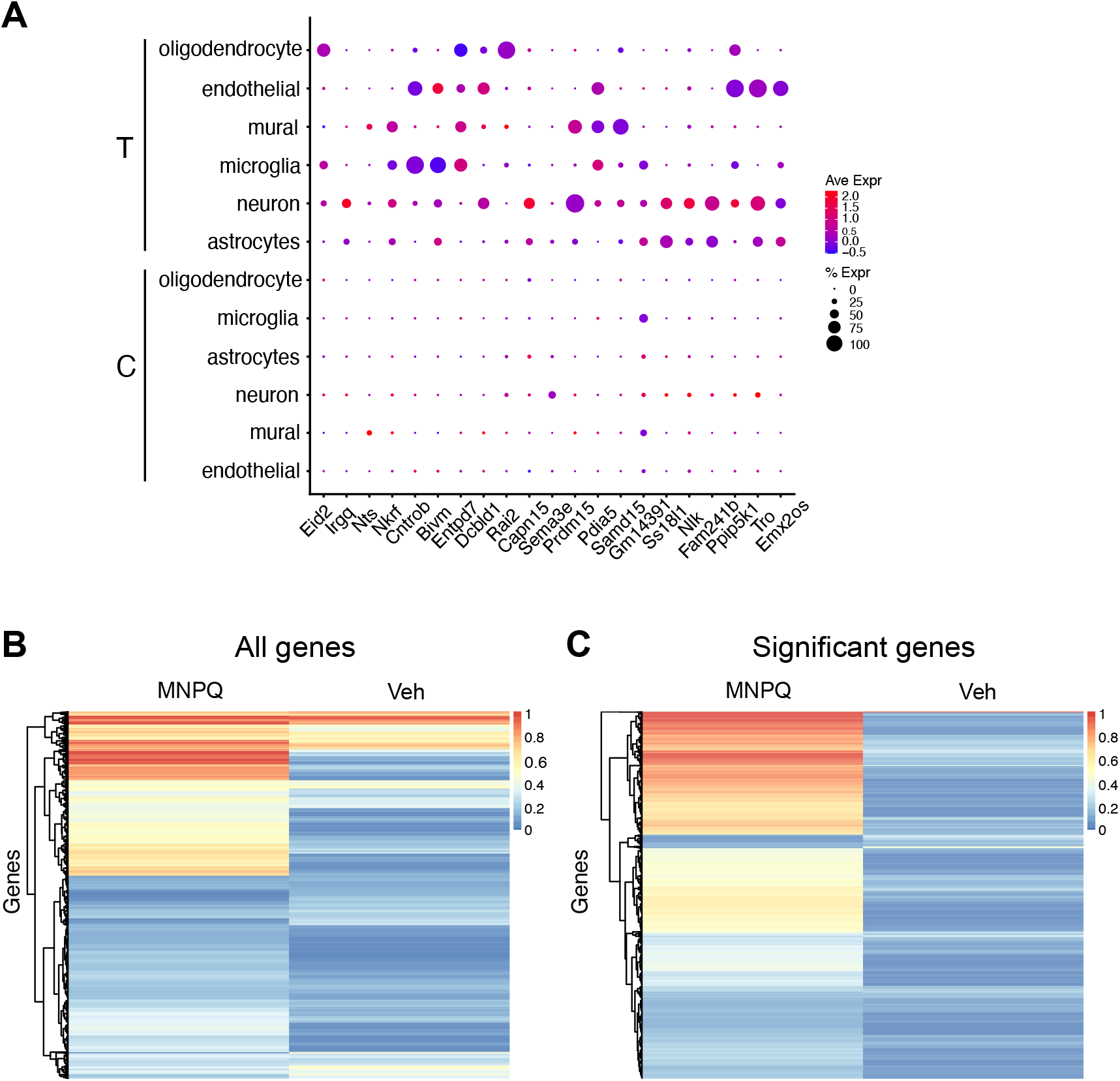
Significant differences in expression between MNPQ and vehicle samples. (**A**) Significant differences in expression between MNPQ and vehicle in each cell type. Expression is generally lower in the treated (T) samples (blue color) compared to controls (C), even though treated samples have higher percentages of expressing cells. Average expression (Ave Expr) denoted by red-blue color scale. Percent of cells expressing gene (% Expr) denoted by dot size. (**B**) Heatmap of percent of cells using all genes from all clusters. Increased cell percentage expressing genes in MNPQ compared to vehicle. (**C**) Heatmap of cell percentage for statistically significant genes.

The increased percentage of expressing cells in the MNPQ mice may represent compensatory induction of low levels of gene expression in normally non-expressing cells. Alternatively, the increased percentage may be an indirect consequence of pesticide-induced cell death selectively destroying non-expressing cells, while sparing any remaining cells with low expression. The genes with low expression may therefore represent a signature of increased cellular resilience to MNPQ.

### Enrichment analysis of significant differentially expressed genes

The 1750 significant differentially expressed genes were analyzed for functional enrichment (Supplementary Table S6). Functions relevant to PD were found to be over-represented using gene ontology (GO) in Enrichr, including regulation of axonogenesis, modulation of chemical synaptic transmission, neuron projection morphogenesis and regulation of AMPA receptor activity (adjusted *P* < 4.7 × 10^−7^) (Kuleshov et al., 2016; The Gene Ontology Consortium et al., 2021).

We also evaluated KEGG pathways in the significant differentially expressed genes (Kanehisa et al., 2007) (Supplementary Table S6). Pathways were enriched in terms relevant to PD, such as GABAergic synapse pathway, dopaminergic synapse and cholinergic synapse (adjusted *P* < 9.4 × 10^−4^). Relevant genes included *Prkacb, Fos, Adcy5* and *Homer1* (Erro et al., 2021; Odumpatta and Arumugam, 2021; Tran et al., 2020; Zhu et al., 2018).

The significant differentially expressed genes were also strongly enriched in terms related to PD in the GeneRIF ARCHS4 predictions of rare diseases (Jimeno-Yepes et al., 2013; Lachmann et al., 2018). These terms included dystonia, neuronal intranuclear inclusion disease, and Parkinson disease juvenile autosomal recessive (adjusted *P* < 3.03 × 10^−24^)., Consistent with the increased risk of PD in older individuals, the significant differentially expressed genes were enriched in downregulated gene expression signatures in the GTEx catalog of aging human brain (20-29 y vs 60-69 y, adjusted *P* = 1.64 × 10^−10^) (The GTEx Consortium, 2020).

### Networks of significant differentially expressed genes

To further dissect genetic pathways in pesticide-induced PD, we used GeneMANIA to analyze the top 100 most significant differentially expressed genes between MNPQ and vehicle treated mice in neurons (Figure 5). GeneMANIA generates hypotheses about gene function by identifying networks of genes with similar roles based on publicly available genomics and proteomics data (Franz et al., 2018).

**Figure 5.**
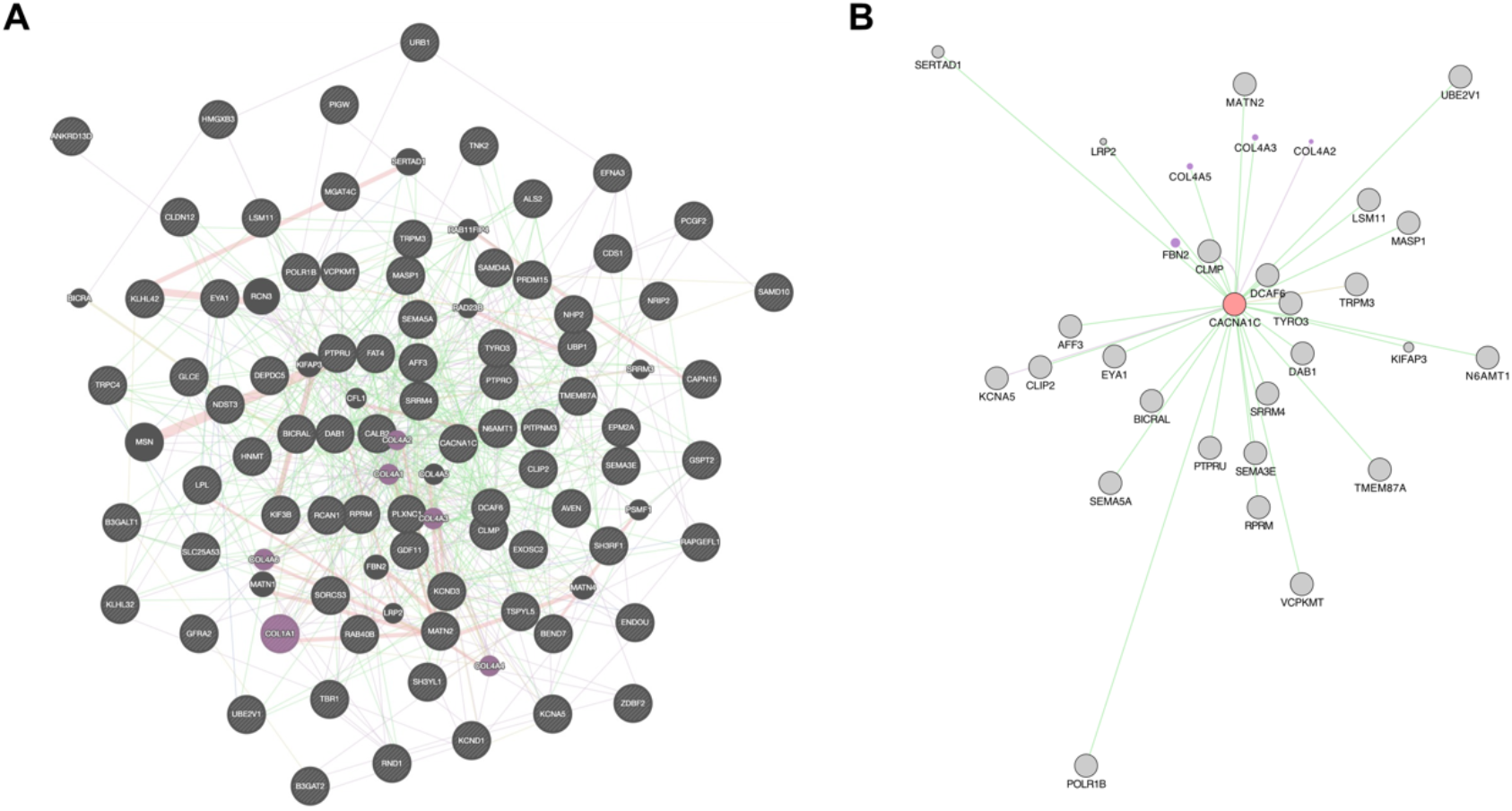
Networks for significant differentially expressed genes in neurons. (**A**) The network is significantly enriched in extracellular matrix genes (purple). The most common interactions are genetic interactions from radiation hybrid genotypes (green). (**B**) Subnetwork featuring Cacna1c.

The most common interactions in the network were due to genetic interactions inferred from radiation hybrid genotypes (Lin et al., 2010). These interactions composed half of the total and were six times more common than the next most frequent, which were from the InterPro classification of protein families (Blum et al., 2021). The network was significantly enriched in terms related to collagen, extracellular matrix and basement membrane (FDR < 8.49 × 10^−4^) (Figure 5A). Other studies have also implicated changes in the extracellular matrix as a result of PD (Aguila et al., 2021; Freitas et al., 2021; Sandor et al., 2017). One hub gene, *Cacna1c*, had 31 interactions (Figure 5B). L-type calcium channels such as *Cacna1c* are implicated in PD, and pharmacological blockade of these channels have been proposed as a potential therapy for the disorder (Kasap and Dwyer, 2021; Liss and Striessnig, 2019; Ritz et al., 2010).

We extended our analysis by using all significant differentially expressed genes to create a network of protein-protein interactions from the InnateDB tool in OmicsNet (Breuer et al., 2013; Zhou and Xia, 2019). InnateDB curates extensive experimentally validated molecular interactions and pathway annotations for both human and mouse. We found that *Nova1* acted as hub gene with 115 linked nodes among the significant differentially expressed genes in all clusters. (Supplementary Figure S6). *Nova1* was significantly more highly expressed in MNPQ neurons than controls (adjusted *P* = 1.03 × 10^−4^). Interestingly, *Nova1* regulates neuron-specific alternative splicing and also binds to a *cis*-regulatory region in the α-synuclein gene, which is linked to PD via both common and rare variants (Jensen et al., 2000; McClymont et al., 2018).

## Discussion

In order to model pesticide-induced PD, we treated C57BL/6J mice with MNPQ or vehicle. Motor effects reminiscent of PD were detected in the MNPQ-treated mice using the pole test. We used scRNA-seq of the SNpc of MNPQ and control mice to understand the molecular signatures of pesticide-induced PD at a cellular level. We expected to capture > 10,000 single cells expressing the transcriptome from most mouse genes. Indeed, using initial filtering conditions, we obtained expression of transcripts from 11252 and 10287 cells for control and experimental samples respectively, which met our expectations (Baquet et al., 2009; Ip et al., 2017). However, when we used stringent filtering criteria to remove dead cells and ambient RNA from the data, only 494 and 468 single cells remained for evaluation. This number of cells is less than ideal, but we decided that exploratory analysis of these pilot data may give some insights into the cellular and molecular mechanisms of pesticide-induced PD.

Unsupervised clustering gave 13 groups at first. However, employing marker genes to assign clusters to specific cell types resulted in a total of 7 clusters. The majority of markers in each cluster were specific except for a few that overlapped between clusters. The overlapping markers may be due to proliferating precursor cells that share transcriptional profiles, even though they are destined to form different cell types (Clarke et al., 2021).

Genes were identified in each cluster with increased expression relevant to dopaminergic neurotransmission and pesticide-induced PD. Examples of cluster specific genes included neuron (*Syt1, Snap25* and *Rtn1*), astrocyte (*Gpr37l1, Pla2g7* and *Prdx6*) and oligodendrocytes (*Ermn, Cldn11* and *Ugt8a*).

Despite the small number of isolated cells, 1750 genes were identified with statistically significant up- and down-regulation between MNPQ and controls. Significant differentially expressed genes were found in all cell clusters, except ependymal. The significant genes also showed some commonality with PD genes identified using genome-wide association, with one gene, *Lrrk2*, also responsible for monogenic PD. The largest number of significant differentially expressed genes were found in neurons, followed by astrocytes and oligodendrocytes. This finding highlights the importance of neuronal support cells in the pathogenesis of pesticide-induced PD and is consistent with previous reports.

A total of 14 genes were significantly differentially expressed in 4 or more cell types in the SNpc, suggesting that cell-agnostic as well as cell-specific pathways play a role in pesticide-induced PD. Functional enrichment analysis of differentially expressed genes using GO highlighted processes such as regulation of axonogenesis, modulation of chemical synaptic transmission, neuron projection morphogenesis and regulation of AMPA receptor activity. In addition, KEGG pathways included GABAergic synapse pathway, dopaminergic synapse and cholinergic synapse.

In a literature database of rare diseases, the significant differentially expressed genes were strongly enriched in terms related to neuronal intranuclear inclusion disease and juvenile PD, highlighting the relationships between pesticide-induced and other causes of PD. The significant differentially expressed genes were also enriched in terms related to gene expression signatures in aging human brain, consistent with PD as mostly a disease of older people.

A network analysis of 100 significant differentially expressed genes in neurons using GeneMANIA underlined the importance of the extracellular matrix in pesticide-induced PD and, together with the relevance of astrocytes and oligodendrocytes, emphasizes that PD is not solely a disorder of neurons. A further network analysis of all differentially expressed genes using InnateDB indicated that *Nova1*, a regulator of alternative splicing, acts as a hub gene and is expressed at higher levels in MNPQ SNpc neurons than controls. This finding draws attention to *Nova1* as a key regulator in PD.

Employing a model of pesticide-induced PD in mice, together with scRNA-seq analysis, we were able to classify different cell types in the SNpc and delineate significant expression differences between pesticide-exposed and control cells. Gene enrichment and network analyses of differentially expressed genes highlighted relevant genetic pathways in the model. Additional study of pesticide-induced PD models using scRNA-seq technology is likely to provide further insights into this environmental cause of PD.

## Acknowledgements

We thank the American Parkinson’s Disease Association Center for Advanced Research (Marie-Françoise Chesselet, Beate Ritz, Principal Investigators) for funding. We also thank the UCLA Semel Institute Neurosciences Genomics Core for sequencing.

