## Supplementary Materials for "Single cell analysis of gene expression in the substantia nigra pars compacta of a pesticide-induced mouse model of Parkinson’s disease"

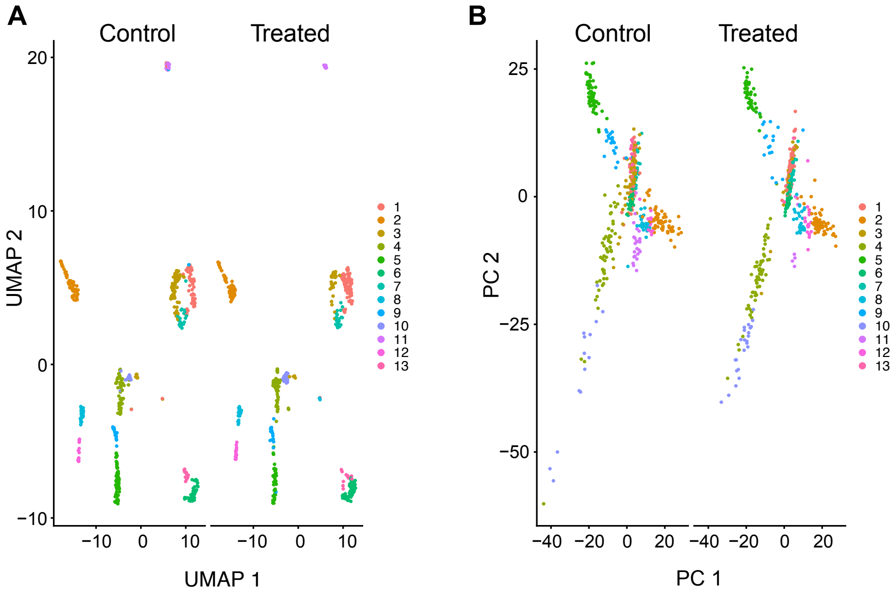


***Supplementary Figure S1.*** *Initial cell clustering of SNpc brain tissues from MNPQ treated and control C57BL/6J mice. (**A**) Uniform manifold approximation and projection (UMAP). (**B**) Principal components (PC) analysis.*

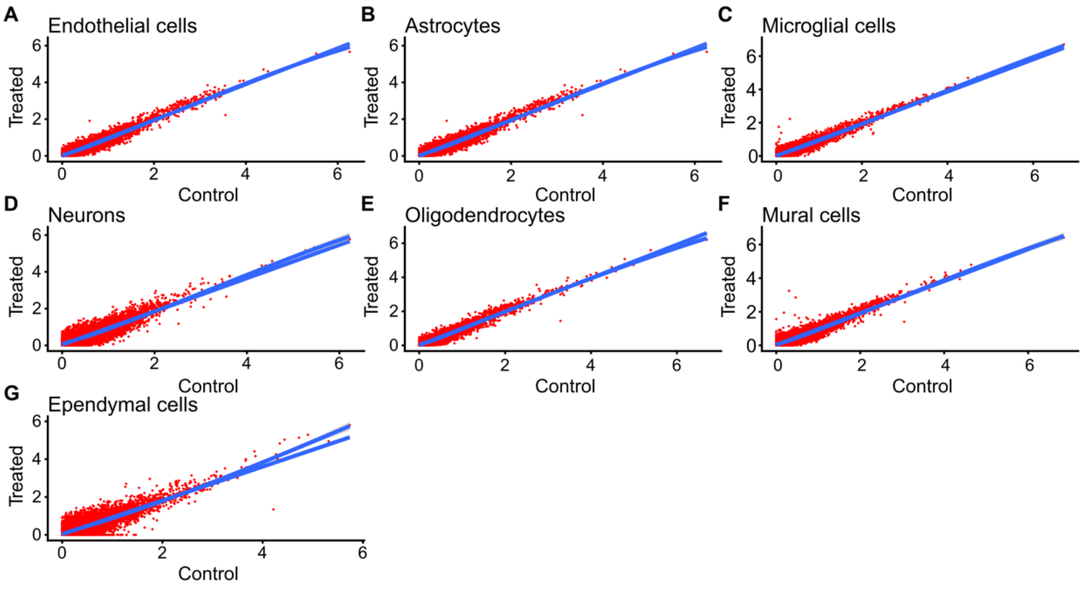


***Supplementary Figure S2.*** *Scatter plots comparing average gene expression between control and MNPQ treated samples for each specific cluster. (**A**) Endothelial cells. (**B**) Astrocytes. (**C**) Microglia. (**D**) Neurons. (**E**) Oligodendrocytes. (**F**) Mural cells. (**G**) Ependymal cells. Markers in each cluster show differential expression. Blue line and shaded blue lines drawn using linear regression and loess methods, respectively.*

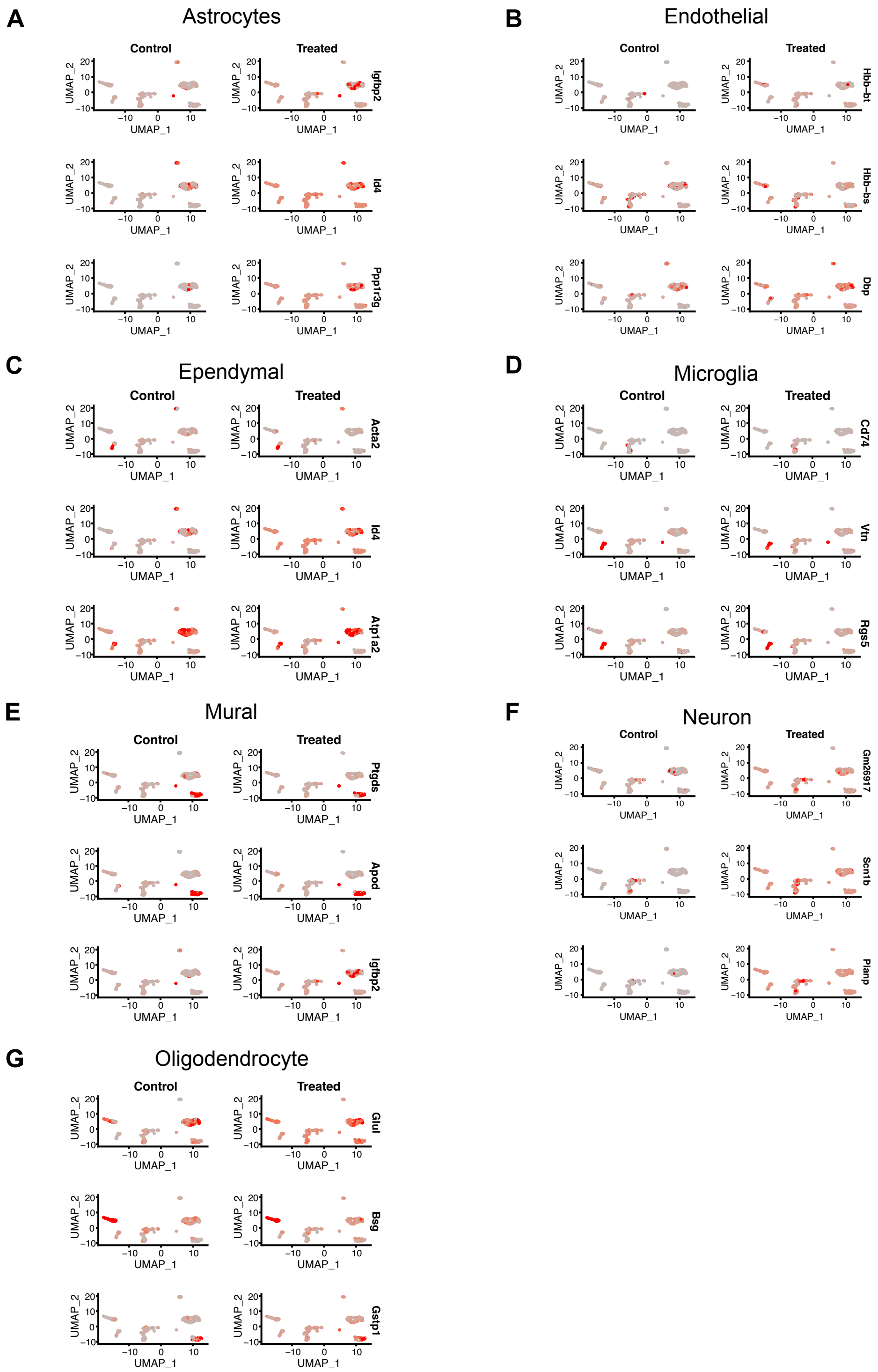


***Supplementary Figure S3.*** *Feature plots for top markers showing largest average differential expression between MNPQ and vehicle in each cluster.* ***(A)*** *Astrocytes.* ***(B)*** *Endothelial cells.* ***(C)*** *Ependymal cells.* ***(D)*** *Microglia.* ***(E)*** *Mural cells.* ***(F)*** *Neurons.* ***(G)*** *Oligodendrocytes. Cluster specific markers showing differences in expression between conditions are evident.*

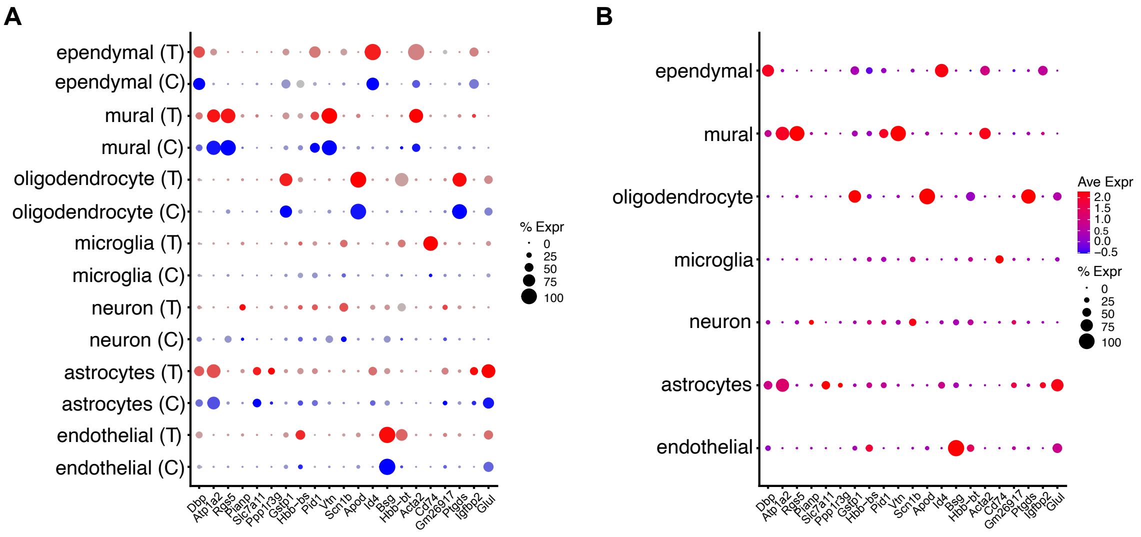


***Supplementary Figure S4.*** *Marker genes from each cluster with large expression differences between MNPQ and control. (**A**) Marker genes for MNPQ treated (T, red-grey expression scale) and control samples (C, blue-grey expression scale). Percent of cells expressing gene (% Expr) denoted by dot size. Marker genes with large MNPQ-vehicle differences in each cluster are Acta2, Id4, Atp1a2, Pid1 (ependymal cells); Ptgds, Apod, Slc7a11 (mural cells); Bsg, Gstp1, Glul (oligodendrocytes); Cd74, Vtn, Rgs5 (microglia); Scn1b, Pianp, Gm26917 (neuron); Igfbp2, Id4, Ppp1r3g, Dbp (astrocytes); Hbb-bt, Dbp, Gstp1 (endothelial cells). (**B**) Same marker genes for combined MNPQ and control samples. Average expression (Ave Expr) denoted by red-blue color scale.*

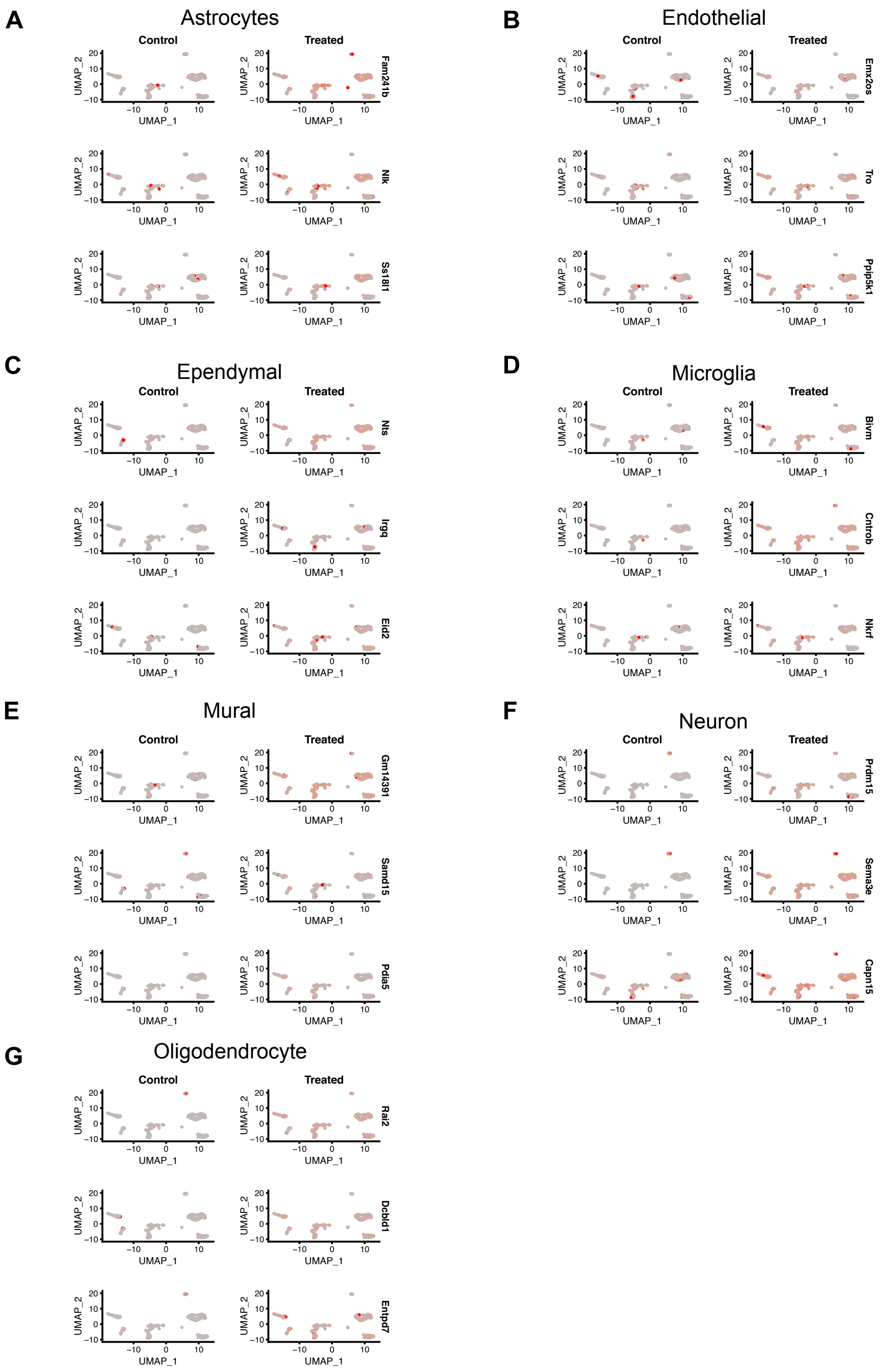


***Supplementary Figure S5.*** *Feature plots of genes in each cluster with significant differential expression between MNPQ and vehicle. (**A**) Astrocytes. (**B**) Endothelial cells. (**C**) Ependymal cells. (**D**) Microglia. (**E**) Mural cells. (**F**) Neurons. (**G**) Oligodendrocytes. Evidence of significant differential expression in most clusters.*

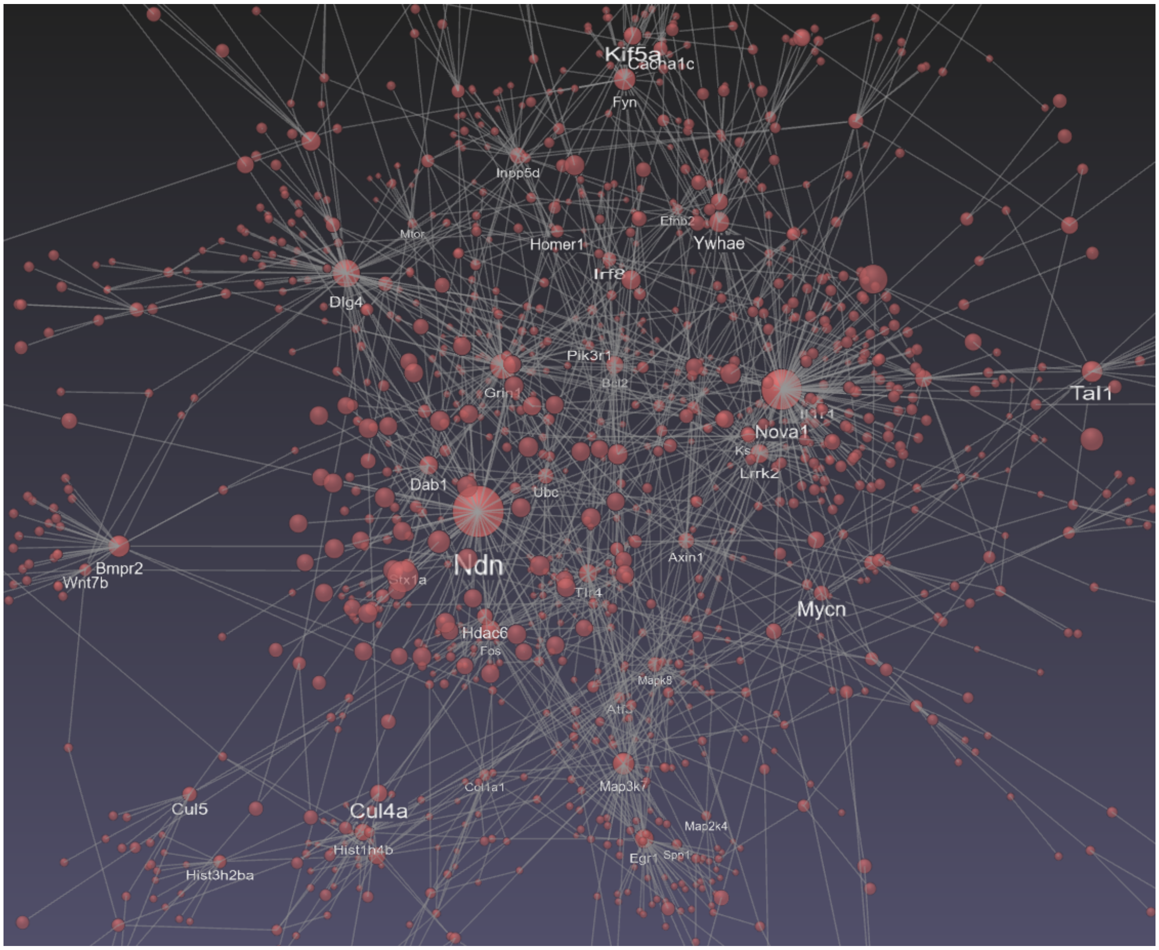


***Supplementary Figure S6.*** *Protein-protein interactions for significant differentially expressed genes. Nova1 acted as a hub gene, connecting with highest number of nodes (115).*
